# Engineering High-Quality Cartilage Microtissues using Hydrocortisone Functionalised Microwells

**DOI:** 10.1101/2022.09.07.507002

**Authors:** Ross Burdis, Gabriela Soares Kronemberger, Daniel J. Kelly

**Affiliations:** Trinity Centre for Biomedical Engineering, Trinity Biomedical Sciences Institute, Trinity College Dublin, Ireland; Department of Mechanical, Manufacturing and Biomedical Engineering, School of Engineering, Trinity College Dublin, Ireland; Advanced Materials and Bioengineering Research Centre (AMBER), Royal College of Surgeons in Ireland and Trinity College Dublin, Ireland; Department of Anatomy and Regenerative Medicine, Royal College of Surgeons in Ireland, Dublin, Ireland

**Keywords:** Mesenchymal Stem/Stromal Cell, Chondrogenesis, Hydrocortisone, Microtissue, Microwell, Biofabrication, Cartilage Tissue Engineering

## Abstract

Engineering clinically-relevant musculoskeletal tissues at a human scale is a considerable challenge. Developmentally-inspired scaffold-free approaches for engineering cartilage tissues have shown great promise in recent years, enabling the generation of highly biomimetic tissues. Despite the relative success of these approaches, the absence of a supporting scaffold or hydrogel creates challenges in the development of large scale tissues. Combining numerous scaled-down tissue units (herein termed *microtissues*) into a larger macrotissue represents a promising strategy to address this challenge. The overall success of such approaches, however, relies on the development of strategies to support the robust and consistent chondrogenic differentiation of clinically relevant cell sources such as mesenchymal stem/stromal cells (MSCs) within microwell arrays to biofabricate numerous microtissues rich in cartilage-specific extracellular matrix components. In this paper, we first describe a simple method to manufacture cartilage microtissues at various scales using novel microwell array stamps. This system allows the rapid and reliable generation of cartilage microtissues, and can be used as a platform to study microtissue phenotype and development. Based on the unexpected discovery that Endothelial Growth Medium (EGM) enhanced MSC aggregation and chondrogenic capacity within the microwell arrays, this work also sought to identify soluble factors within the media capable of supporting robust differentiation using heterogeneous MSC populations. Hydrocortisone was found to be the key factor within EGM that enhanced the chondrogenic capacity of MSCs within these microwell arrays. This strategy represents a promising means of generating large numbers of high-quality, scaffold-free cartilage microtissues for diverse biofabrication applications.

## Introduction

Unlike traditional scaffold or hydrogel-based tissue engineering strategies, scaffold-free approaches are inherently reliant on the cell’s own capacity to generate the bulk of the tissue/construct through the deposition of extracellular matrix (ECM). Although this follows a developmentally inspired paradigm and facilitates the generation of biomimetic *in vitro* cartilage tissues [1–10], in the absence of an interstitial ‘bulking’ scaffold or hydrogel material, creating tissues of scale can be challenging. Scaling down scaffold-free tissue units can support more robust differentiation, alleviate diffusion gradients, and ultimately improve matrix deposition [11–14]. Despite the numerous biological benefits associated with scaled-down 3D scaffold-free strategies [1,15,16], microtissues do not yet represent an idealised solution whereby robust ECM biosynthesis is guaranteed. A number of different stem/progenitor cell sources, including articular chondrocytes [13,17], mesenchymal stem/stromal cells (MSCs) [18–21], periosteal derived stem cells [12], and induced pluripotent stem cell (iPSC) derived cells [22], have been used for the biofabrication of cartilage microtissues. Multiple methods for forming multicellular spheroids/microtissues have been described [23], leveraging various non-adherent polymers [21,24–27] and hydrogels [12–14,28–33] as substrate materials. At present, cartilage microtissues are typically formed in a medium-to-high throughput manner using microwell moulds, whereby stem/progenitor cells are collected in the bottom of a non-adherent well and undergo cellular self-assembly/self-organisation to form a cellular spheroid. Under the appropriate exogenous soluble cues, the cells within these aggregates can be differentiated and begin to deposit a tissue-specific extracellular matrix (ECM), generating a microtissue. An ideal platform for generating microtissues for biofabrication can be defined as a scalable process, capable of supporting the development of standardised spheroids with defined shape, size and phenotype which can be used as part of a subsequent biofabrication strategy, such as bioprinting [34]. Closely coupled with the suitability of the method of forming microtissue building-blocks, is the quality of the microtissue formed. The chosen platform for generating microtissues should support key processes such as differentiation, phenotype commitment/maintenance, and the capacity for subsequent tissue fusion [34]. Evaluation of microtissue quality (ECM composition, cellular phenotype, and functionality) can be carried out using biochemical, histological, gene expression, and fusion assays.

Cellular heterogeneity, particularly with human MSCs, can result in poor chondrogenesis and impact the richness of any cartilage ECM generated *in vitro* [35]. Such donor-to-donor variation is well documented throughout the literature and has been shown to directly affect the *in vivo* performance of engineered cartilages [36]. Therefore, engineering numerous microtissues, rich in cartilage-specific ECM, in a practical and economical manner is challenging using MSC populations considering their inherently variable chondrogenic capacity. The increased interest in the use of cellular spheroids, microtissues, and/or organoids as biological building-blocks for engineering functional osteochondral tissues/organs demands the development of strategies ideally suited to the biofabrication of large numbers of homogeneous and phenotypically defined microtissues. Engineering large numbers of such microtissues using clinically relevant cell sources requires the careful consideration of culture conditions that regulate key outcomes such as microtissue phenotype (e.g. cell types and specific growth factors), quality (e.g. mitigating diffusion gradients and nutrient limitations) and size (e.g. cell numbers) [37]. In the context of cartilage and osteochondral tissue engineering, early research in this area has focused on the use of undifferentiated MSC aggregates [38,39]. These immature aggregates do not mimic the complex native ECM, which may explain why they fail to promote the regeneration of hyaline cartilage when implanted into pre-clinical models for chondral/osteochondral defects [40,41]. Therefore new approaches for engineering high quality cartilage microtissues at scale using clinically relevant cell sources are required.

With this in mind, generating cartilage and osteochondral tissues of scale using microtissue/aggregate engineering will require the high-throughput production of consistently high-quality cartilage microtissues. In particular, methods for consistently engineering quality cartilage microtissues from diverse donors with different chondrogenic capacity is required. Without well-defined markers for identifying MSCs within primary isolations, many tissue engineers use uncharacterised cohorts of cells for generating tissues, which in turn can reduce the reliability and quality of the engineered cartilages. The identification of a simple method for improving the chondrogenic capacity of uncharacterised MSC populations, isolated from bone marrow, could help to limit the variability seen in cartilage tissue engineering. Specifically, in the context of cartilage microtissue/aggregate engineering, the identification of protocols combatable with microwell platforms typically used in the biofabrication of such microtissues are required. Ultimately, these high-quality cartilage microtissues can be used as building blocks to more efficiently engineer cartilage and osteochondral tissues of scale. In this paper we first describe the design of two microwell arrays that can be used to directly pattern a hydrogel into an ideal platform for engineering cartilage microtissues. We demonstrate the capacity to consistently form spherical cell aggregates within both the medium- and high-throughput microwell systems, generating cartilage microtissues of different maturities/phenotypes. Based on a serendipitous observation that Endothelial Growth Medium (EGM) enhanced MSC aggregation and chondrogenic capacity in bone-marrow derived MSCs (BMSCs), this study also sought to elucidate the driving factor(s) supporting such differentiation within EGM. Ultimately, our aim was to improve upon current chondrogenic culture regimes and create a novel platform for engineering high-quality, scaffold-free cartilage microtissues at scale.

## Materials & Methods

### Media Formulations

#### Expansion Medium “XPAN”

XPAN is composed of high glucose Dulbecco’s modified eagle’s medium (hgDMEM) GlutaMAX supplemented with 10 % v/v FBS, 100 U/mL penicillin, 100 μg/mL streptomycin (all Gibco, Biosciences, Dublin, Ireland) and 5 ng/mL FGF2 (Prospect Bio).

#### Endothelial Growth Medium (EGM)

EGM is composed of Endothelial Cell Basal Medium-2 (EBM) (Lonza) supplemented with MV Microvascular Endothelial Cell Growth Medium-2 BulletKit™ (Lonza). As the concentration of the supplements added to EBM are proprietary information, the concentrations for each are given as a %v/v. Fetal bovine serum (FBS) was added at 5 %, Hydrocortisone (Hydro) was added at 0.04 %, human FGF-2 (FGF) was added at 0.4 %, vascular endothelial growth factor (VEGF), recombinant human long R3 insulin like growth factor 1 (IGF), ascorbic acid (AA), and human epidermal growth factor (EGF) were all added at 0.1 %. Finally, gentamicin sulfate-Amphotericin (GA-1000) was added at 0.1 %.

#### Chondrogenic Differentiation Medium (CDM)

hgDMEM GlutaMAX supplemented with 100 U/mL penicillin, 100 μg/mL streptomycin (both Gibco), 100 μg/mL sodium pyruvate, 40 μg/mL L-proline, 50 μg/mL L-ascorbic acid-2-phosphate, 4.7 μg/mL linoleic acid, 1.5 mg/mL bovine serum albumin, 1 X insulin–transferrin–selenium (ITS), 100 nM dexamethasone (all from Sigma), 2.5 μg/mL amphotericin B and 10 ng/mL of human transforming growth factor-β3 (TGF-β) (Peprotech, UK).

*Hypertrophic Differentiation Medium* (*HYP*) was composed of hgDMEM GlutaMAX supplemented with 100 U/ml penicillin, 100 μg/mL streptomycin (both Gibco), 1 × ITS, 4.7 μg/ml linoleic acid, 50 nM thyroxine, 100 nM dexamethasone, 250 μM ascorbic acid, 7 mM β-glycerophosphate and 2.5 μg/mL amphotericin B (all from Sigma).

### Bone Marrow Mesenchymal Stem/Stromal Cell (BMSC) Isolation

#### Goat BMSC (gBMSC) isolation

gBMSCs were harvested under sterile conditions from the sternum of skeletally mature, female, Saanen goats. Briefly, excised bone marrow was dissected into small pieces using a scalpel. The marrow pieces were then gently rotated for 5 min in XPAN to help liberate the cellular components. The culture medium was then aspirated and passed through a 70 μm cell sieve prior to counting and plating at a density of 57 × 10^3^ cells/cm^2^ and expanded under hypoxic conditions (37 °C in a humidified atmosphere with 5 % CO_2_ and 5 % O_2_) for chondrogenic differentiation. Following colony formation, gBMSCs were trypsinised using 0.25 % (w/v) Trypsin Ethylenediaminetetraacetic acid (EDTA). gBMSCs for microtissues were expanded from an initial density of 5000 cells/cm^2^ in XPAN medium under physioxic conditions until P3.

#### Human BMSC (hBMSC) isolation

hBMSCs were isolated from unprocessed human bone marrow (Lonza) on the basis of plastic adherence. Briefly, unprocessed bone marrow was plated at 2.5 × 10^5^ cells/cm^2^ (estimated approx. 4000 - 5000 MSCs/cm^2^) in XPAN medium and expanded under physiological oxygen conditions (37 °C in a humidified atmosphere with 5 % CO_2_ and 5 % O_2_). Following colony formation, hBMSCs were trypsinised using 0.25 % (w/v) Trypsin Ethylenediaminetetraacetic acid (EDTA), hBMSCs were expanded in XPAN under physioxic conditions (5 % O_2_) and aggregated into pellets at P3.

### Microwell Platform Design, Fabrication and Validation

Microwell stamps were designed using Soldiworks CAD software. A summary of the dimensions of the positive microwell stamps can be found in Figure 1A. Both the medium- and high-throughput microwell arrays were designed to avoid flat sections between adjacent microwells. The medium-throughput wells were designed to maintain discrete microtissues within individual microwells from extended culture periods, and as such had a relatively deep well. In contrast, the microwells in the high-throughput system were designed to maximise the number of microwells per macro-well, making each well considerable smaller in dimension. The base of the high-throughput wells was designed to be flat to maintain print fidelity as creating a curved or pointed base would require dimensions that exceeded the printer’s resolution (Both in x,y directions and laser spot size). Both microwell stamps were fabricated using a Form 3 stereolithography (SLA) printer and the high-temperature resin (V2) (both Formlabs, Massachusetts, United States). Prior to printing, a STL file for the part was prepared using Preform 2.16.0 software (Formlabs, Massachusetts, United States), setting a 0.025 μm layer height defined the resolution of the print. Completed parts were washed in propan-2-ol (Sigma Aldrich) to clear any uncured resin, following which they were exposed to UV light (405 nm, 9.1 W) (Form cure, Formlabs, Massachusetts, United States) for 120 min at 80°C to ensure complete crosslinking. Before use, stamps were autoclave sterilised. Hydrogel microwells were moulded using the same procedure as previously described [10]. Briefly, under sterile conditions, 4 % (w/v) molten agarose was patterned using the microwell stamps within the wells of a 6 well-plate. Once cooled, the stamps were removed and the agarose microwells soaked overnight in an appropriate media type before cell seeding.

**Figure 1.**
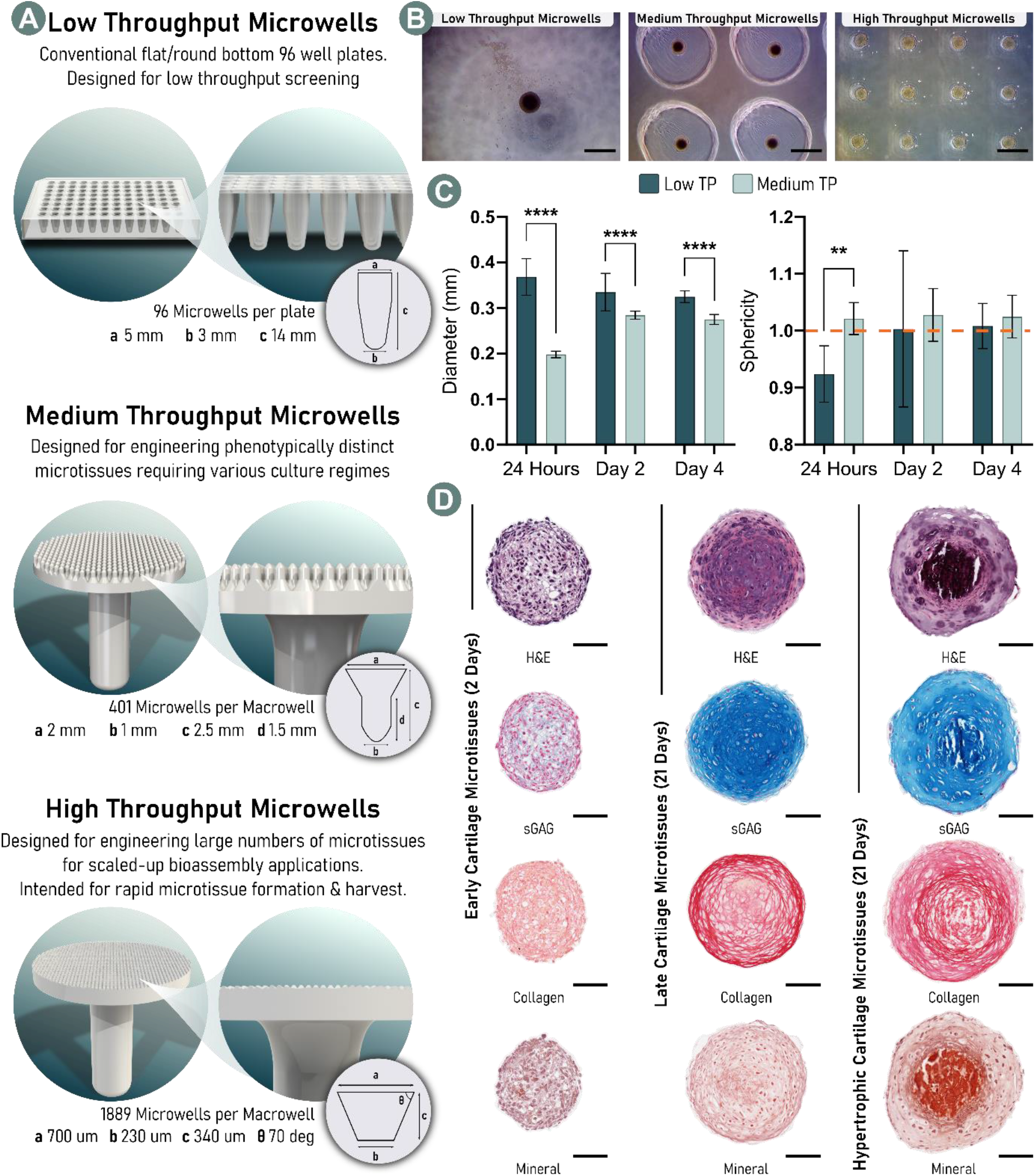
Microwell platform design and validation. A) Schematic representation of three platforms for generating spherical cartilage microtissues. Our medium- and high-throughput systems have been designed to directly pattern a hydrogel substrate within a conventional 6 well-plate. B) Microscopic images of spherical cellular aggregates formed after 2 days within each platform (Scale Bar = 500 μm). C) Quantification of microtissue diameter and sphericity (orange line indicates a perfect sphere) after 24 hours, 2 days, and 4 days of cultivation within the low- and medium-throughput systems. Significant differences were tested using a Šídák’s multiple comparisons test, ordinary two-way ANOVA, where; ** denotes p < 0.01 and **** denotes p < 0.0001, (N = 10, Mean ± SD). D) Histological analysis of phenotypically distinct cartilage microtissues displaying markers of early-, late-, and hypertrophic-cartilage (Scale Bar = 100 μm).

To validate the capacity of the hydrogel microwells to from spherical cellular aggregates and cartilage microtissues, an appropriate cell suspensions (gBMSC) was pipetted into the macrowells to achieve a final density of 4 × 10^3^ cells/microwell. Well plates were then centrifuged at 700 x g for 5 minutes, and returned to physioxic conditions (37 °C in a humidified atmosphere with 5 % CO_2_ and 5 % O_2_) overnight to allow aggregation to occur (^~^18 hours). The following day, media was exchanged to induce chondrogenesis (CDM) and changed every two days until the end point. For hypertrophic cartilage microtissues, after 14 days of chondrogenic cultivation (CDM) HYP media was used for a further 7 days to induce mineralisation of the cartilage microtissues.

### Experimental Design

For all studies, microwells were seeded at a density that results in 4 × 10^3^ cells/microtissue. Cell expansion and cartilage microtissue cultivation took place at physiological oxygen conditions (37 °C in a humidified atmosphere with 5 % CO_2_ and 5 % O_2_). With the exception of the first 24 hours, media was exchanged every 2 days. Initial investigation using gBMSCs involved 21 Days of chondrogenic culture. All studies using hMSCs were carried out over 7 days of chondrogenic cultivation.

#### Control Group

Here, cartilage microtissues are formed by seeding bMSCs into the microwells in XPAN. The following day, the XPAN medium is carefully aspirated from the wells and replaced with CDM.

#### EGM Group

EGM was used to soak the agarose hydrogel microwells overnight prior to seeding. As such, cells seeded into the microwells in the ‘EGM’ group were never directly exposed to EGM. The seeding procedure and following chondrogenic culture was identical to the control group.

#### Media Component Isolation Groups

By means of determining the prominent factor within EGM that aided in chondrogenesis, a screening study was undertaken. Each of the supplements listed in §2.1 for EGM formulation were added at the correct concentration (%v/v) to both XPAN during pre-soaking and seeding, as well as CDM during differentiation culture. Chondrogenic and experimental EGM supplements were added to the basal media of CDM fresh prior to media exchange. Additionally, blends of XPAN/CDM and EGM were used. In these groups EGM was supplemented as a 1× or 2× formulation and then mixed 50/50 with either XPAN, for soaking and seeding, or with CDM (2×) for chondrogenic differentiation. Summaries of the experimental groups and media compositions can be found in supplementary figure 1 and supplementary table 1.

### Histological Analysis

Samples were fixed using 4 % paraformaldehyde (PFA) solution overnight at 4 °C. After fixation, samples were dehydrated in a graded series of ethanol solutions (70 % - 100 %), cleared in xylene, and embedded in paraffin wax (all Sigma-Alrich). Prior to staining tissue sections (5 μm) were rehydrated. Sections were stained with haematoxylin and eosin (H&E), 1 % (w/v) alcian blue 8GX in 0.1 M hydrochloric acid (HCL) (AB) to visualise sulphated glycosaminoglycan (sGAG) content and counter-stained with 0.1 % (w/v) nuclear fast red to determine cellular distribution, 0.1 % (w/v) picrosirius red (PSR) to visualise collagen deposition, and 1 % (w/v) alizarin red (pH 4.1) to determine mineral deposition *via* calcium staining (all from Sigma-Aldrich). Stained sections were imaged using an Aperio ScanScope slide scanner.

### Quantitative Biochemical Analysis

Samples were washed in PBS after retrieval and the number of microtissues within each technical replicate counted prior to digestion. A papain enzyme solution, 3.88 U/mL of papain enzyme in 100mM sodium phosphate buffer/5mM Na2EDTA/10mM Lcysteine, pH 6.5 (all from Sigma–Aldrich), was used to digest the samples at 60 °C for 18 hours. DNA content was quantified immediately after digestion using Quant-iT™ PicoGreen ^®^ dsDNA Reagent and Kit (Molecular Probes, Biosciences). The amount of sGAG was determined using the dimethylmethylene blue dye-binding assay (Blyscan, Biocolor Ltd., Northern Ireland), with a chondroitin sulphate standard read using the Synergy HT multi-detection micro-plate reader (BioTek Instruments, Inc) with a wavelength set to 656 nm. Total collagen content was determined using a chloramine-T assay [42] to measure the hydroxyproline content and calculated collagen content using a hydroxyproline-to-collagen ratio of 1:7.69. Briefly, samples were mixed with 38 % HCL (Sigma) and incubated at 110 °C for 18 hours to allow hydrolysis to occur. Samples were subsequently dried in a fume hood and the sediment reconstituted in ultra-pure H2O. 2.82 % (w/v) Chloramine T and 0.05 % (w/v) 4-(Dimethylamino) benzaldehyde (both Sigma) were added and the hydroxyproline content quantified with a trans-4-Hydroxy-L-proline (Fluka analytical) standard using a Synergy HT multi-detection micro-plate reader at a wavelength of 570 nm (BioTek Instruments, Inc).

### Image Quantification & Statistical Analysis

Diameter measurement of growing microtissues were taken from microscope images (4×) using ImageJ software. Statistical analysis was performed using GraphPad Prism software (GraphPad Software, CA, USA). Analysis of differences between two groups at one timepoint was done using a standard two-tailed t-test. For two groups over multiple time-points a one-way analysis of variance (ANOVA) was performed. Numerical and graphical results are presented as mean ± standard deviation unless stated otherwise. Significance was determined when p <0.05.

## Results

### Microwell Platforms for Microtissue Biofabrication

We developed two microwell platforms ideally suited to engineer numerous cartilage microtissues and compared these to standard 96 well plates (Figure 1A). Both the medium- and high-throughput microwell stamps were designed to directly pattern an agarose hydrogel within a conventional 6 well-plate [10]. The medium-throughput stamp generates 401 round bottom microwells with a sufficient well depth to maintain discrete microtissues within individual microwells for extended culture periods allowing for numerous media changes. Such platforms can be used for the engineering of cartilage microtissues with various phenotypes (Figure 1D) and/or investigating microtissue development under various culture conditions. Although all platforms support the formation of spherical cell aggregates (Figure 1B), we demonstrated that our custom microwell platforms result in a more rapid and reliable spheroid formation when compared to a conventional 96 well plate (Figure 1C). The medium-throughput hydrogel microwell was hereon in used as the platform for investigating novel culture conditions for supporting enhanced chondrogenesis.

### Endothelial Growth Media (EGM) Treatment Enhances Aggregation and Chondrogenesis of BMSCs

Soaking agarose hydrogel microwells with EGM prior to cell seeding appeared to have a rapid and potent effect on the self-organisation of gBMSCs into a cellular spheroid. By day 2, gBMSC aggregates were significantly larger, and microscopically appeared to include all of the cells which had been seeded into the individual microwells (Figure 2Ai). In contrast, gBMSC aggregates generated within XPAN soaked microwells had a smaller average diameter, with a large numbers of cells not coalescing within the spheroid, instead appearing at the bottom of the microwell. These significant differences in size were maintained throughout the culture period, resulting in a final average microtissue diameter of 0.403 ± 0.03 μm and 0.311 ± 0.026 μm for EGM and XPAN soaked microwells respectively (Figure 2Aii). Histologically, both microtissue cohorts exhibited canonical markers for chondrogenic differentiation, with positive matrix staining for sGAG and collagen deposition. In the EGM pre-soak group, the intensity of the staining indicated a richer cartilaginous ECM. Neither group stained positive for calcium deposition, providing evidence that the cartilage has not yet progressed towards a mature hypertrophic phenotype (Figure 2B). Biochemical evaluation demonstrated that there were significantly higher levels of DNA, sGAG, and collagen per microtissue when the microwells were soaked with EGM compared to XPAN. Additionally, the levels of sGAG and collagen deposited, normalised to DNA content, demonstrated that EGM treatment resulted in a higher biosynthetic output at a cellular level (Figure 2C). Collectively, these results indicated that EGM treatment resulted in the generation of larger, more cellular microtissues containing higher levels of cartilage-specific ECM components. Moreover, the cells within the microtissue demonstrated a higher synthetic output compared to those undergoing a traditional chondrogenic culture regime.

**Figure 2.**
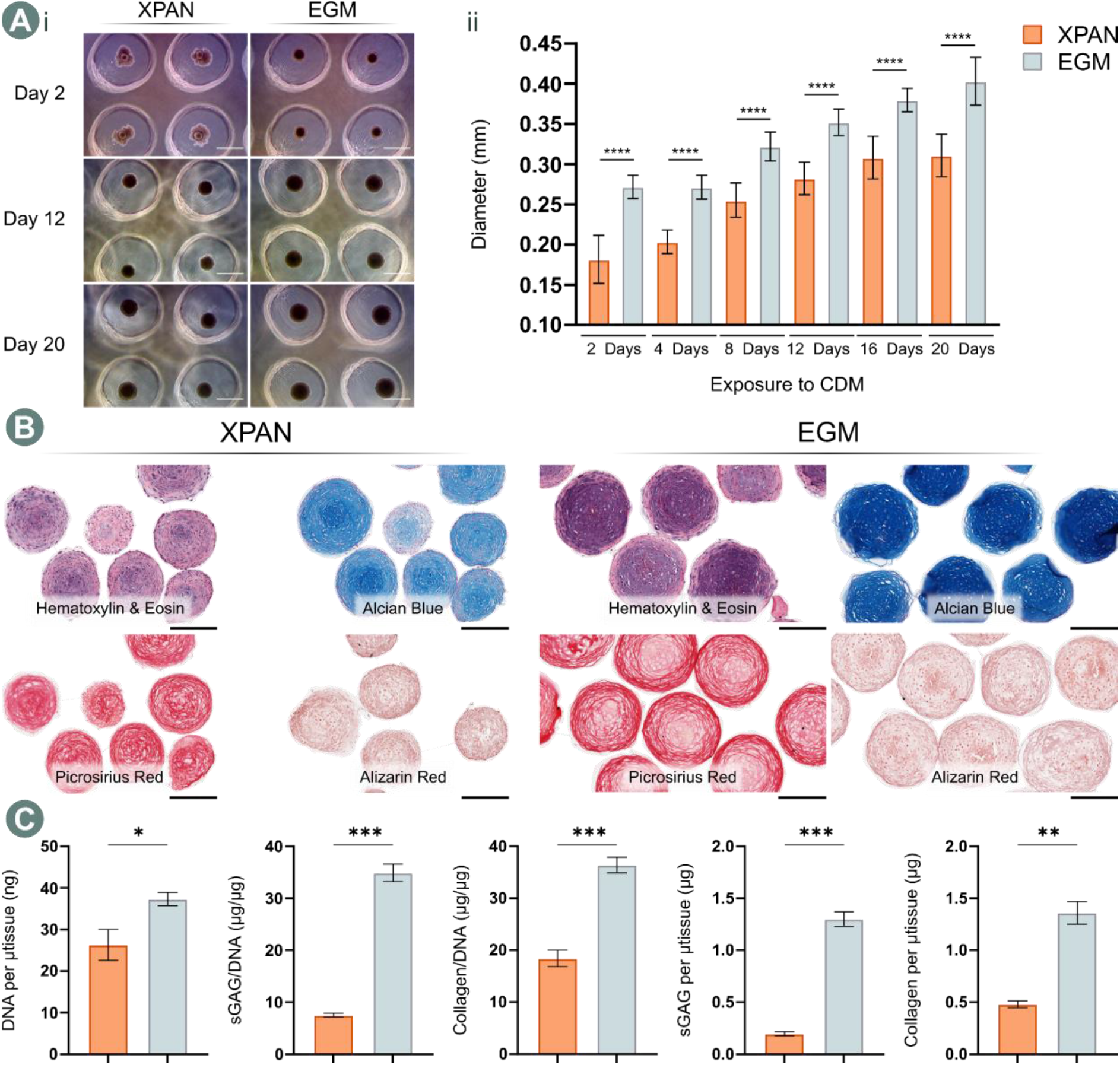
Soaking hydrogel microwells with EGM results in a richer matrix within cartilage microtissues. Ai) Microscopic images at days 2, 12, and 20 during chondrogenic culture (Scale bar = 500 μm), and quantification of diameter (ii) as a non-destructive metric for microtissue development. **** denotes significance when tested using a Šídák’s multiple comparisons test, two-way ANOVA, where p < 0.0001, (N = 20, Mean ± SD). B) Histological evaluation of cartilage microtissues after 21 days of chondrogenic culture (Scale bar = 200 μm). C) Biochemical quantification of the cartilage microtissues after 21 days. * denotes significance using a two-tailed, unpaired Welch’s t-test, where; * indicates p < 0.05, ** indicates p <0.01, and *** indicates p < 0.001 (N = 3, Mean ± SD)

### Hydrocortisone Supports Enhanced Chondrogenesis in Human BMSCs

EGM contains multiple factors that potentially enhance chondrogenesis of BMSCs in this microtissue model (Supplementary table 1). This motivated an investigation to identify the predominant driving factors within the EGM. Additionally, to improve its clinical relevance, this empirical study was undertaken using hBMSCs. To this end, hBMSCs within the microwell system were cultured in media supplemented with each factor used within the EGM formulation. These factors were added to both the XPAN used during soak loading and seeding, as well as to CDM during chondrogenic induction. A typical chondrogenic culture regime ‘control’ was also carried out, as well as EGM soak loading (here termed ‘EGM’), which was identical to the protocol shown to be effective in animal derived BMSCs. After 7 days of *in vitro* chondrogenesis, differences in microtissue size were apparent microscopically (Figure 3B). Cellular arrangement also appeared to vary within the microtissues depending on which additional EGM supplement was provided (Figure 3A). Histologically, archetypal cartilage spheroids were seen in the control, EGM, and hydrocortisone (Hydro) groups. Although other supplements did not entirely suppress chondrogenesis, with sGAG and collagen deposition detected in all groups at varying levels, they did result in condensed, highly cellular and atypical cartilage spheroids. When compared to a standard chondrogenic culture regime (control), supplementation with hydrocortisone resulted in significantly higher levels of DNA per microtissue, as well as a higher deposition of sGAG per cell (Figure 3C). Although EGM soaking did not significantly influence the DNA levels within the microtissues in hBMSCs, its effect on biosynthetic output did mirror observations made previously with gBMSCs, whereby pre-soaking with EGM resulted in a significantly richer cartilaginous ECM profile compared to standard chondrogenic culture conditions (control).

**Figure 3.**
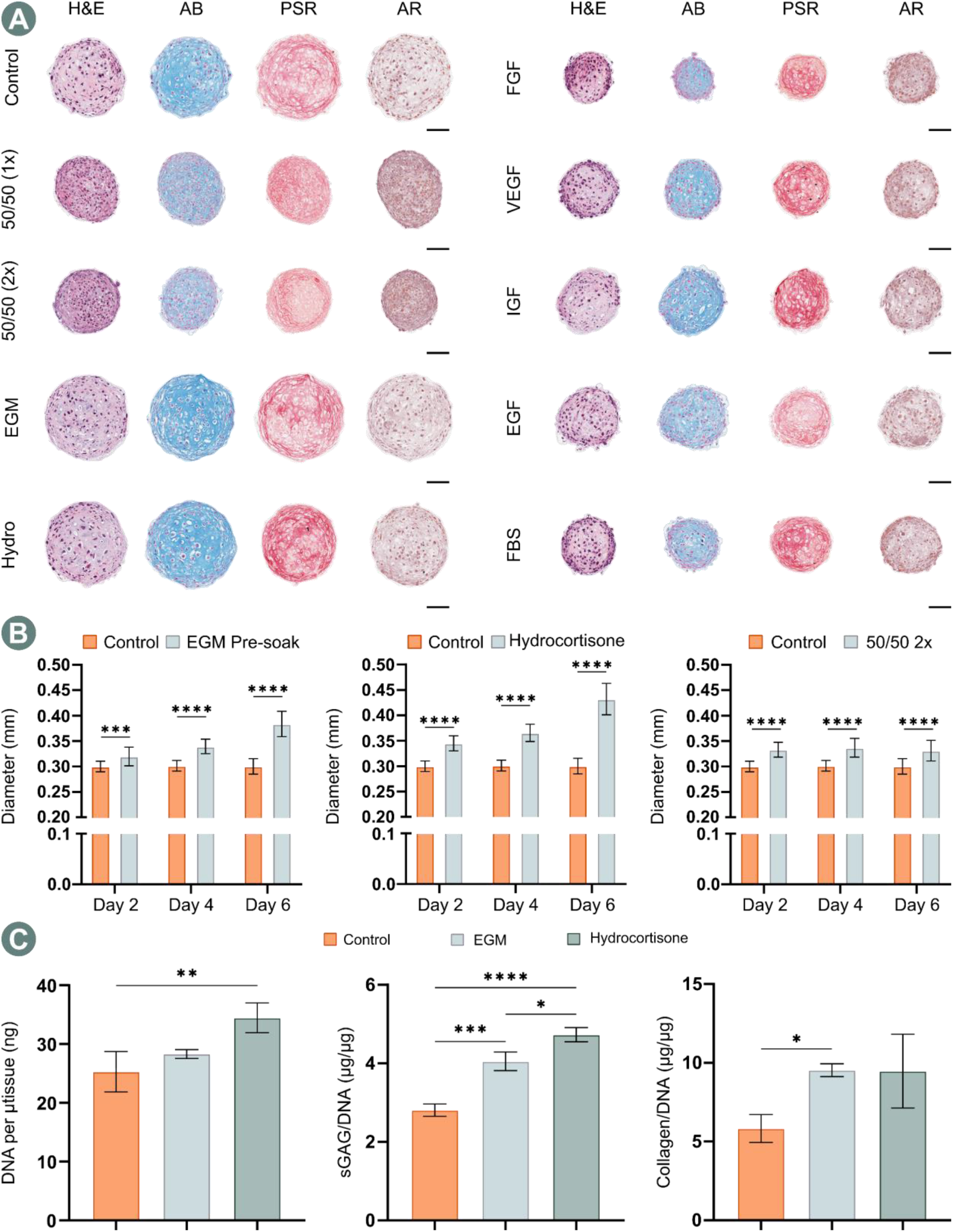
Hydrocortisone is the driving factor in EGM that improved chondrogenesis. A) Histological panel of microtissues, representing each EGM supplement, after 7 days of chondrogenic culture (Scale Bar = 100 μm). B) Diameter measurements during culture, * denotes significance when tested using a Šídák’s multiple comparisons test, two-way ANOVA, where; *** indicates p < 0.001 and **** indicates p < 0.0001, (N = 20, Mean ± SD). C) Biochemical quantification of the cartilage microtissues after 7 days of chondrogenic culture. Hydrocortisone treatment compared to typical chondrogenic conditions (control) and positive control group (EGM) demonstrated a significant increase in DNA content and sGAG deposition. * denotes significance using an Ordinary One-way ANOVA with a Tukey’s multiple comparisons test, where; * indicates p < 0.05, ** indicates p < 0.01, and *** indicates p < 0.001, and **** indicates p < 0.0001 (N = 3, Mean ± SD).

To confirm the effect of EGM and hydrocortisone treatment on hMSC chondrogenesis in this microtissue system, the key groups from the above experiment were repeated using cells isolated from a different human donor. Diameter measurements taken during the 7 days *in vitro* revealed similar responses in terms of microtissue growth in both experimental groups (Figure 4A). Although microtissues within these groups remained significantly larger than those under conventional chondrogenic conditions, unlike in the previous study, the diameter of microtissues in the control group also increased over the 7 days. Histologically, all groups supported robust chondrogenic differentiation and the deposition of cartilage specific ECM components (Figure 4B). Biochemical quantification of the cartilage microtissues indicated that significantly higher levels of sGAG/DNA and sGAG/microtissue could be achieved using the EGM soak loading and hydrocortisone treatments respectively. Despite this, there was no significant benefit in terms of collagen deposition for either experimental group. Interestingly, the baseline chondrogenic capacity of the donor investigated within this study appeared far superior to that of the previous hBMSCs. Under standard chondrogenic conditions (control), the sGAG/DNA was 5.50 ± 0.175, compared to 2.81 ± 0.236 for the previous donor. The difference in collagen deposition per cell was more pronounced, with 25.9 ± 1.13 collagen/DNA for this donor versus 5.83 ± 0.887 collagen/DNA for the previous donor. This suggests that the beneficial effects of such treatments are more pronounced when the baseline levels of chondrogenesis are relatively low.

**Figure 4.**
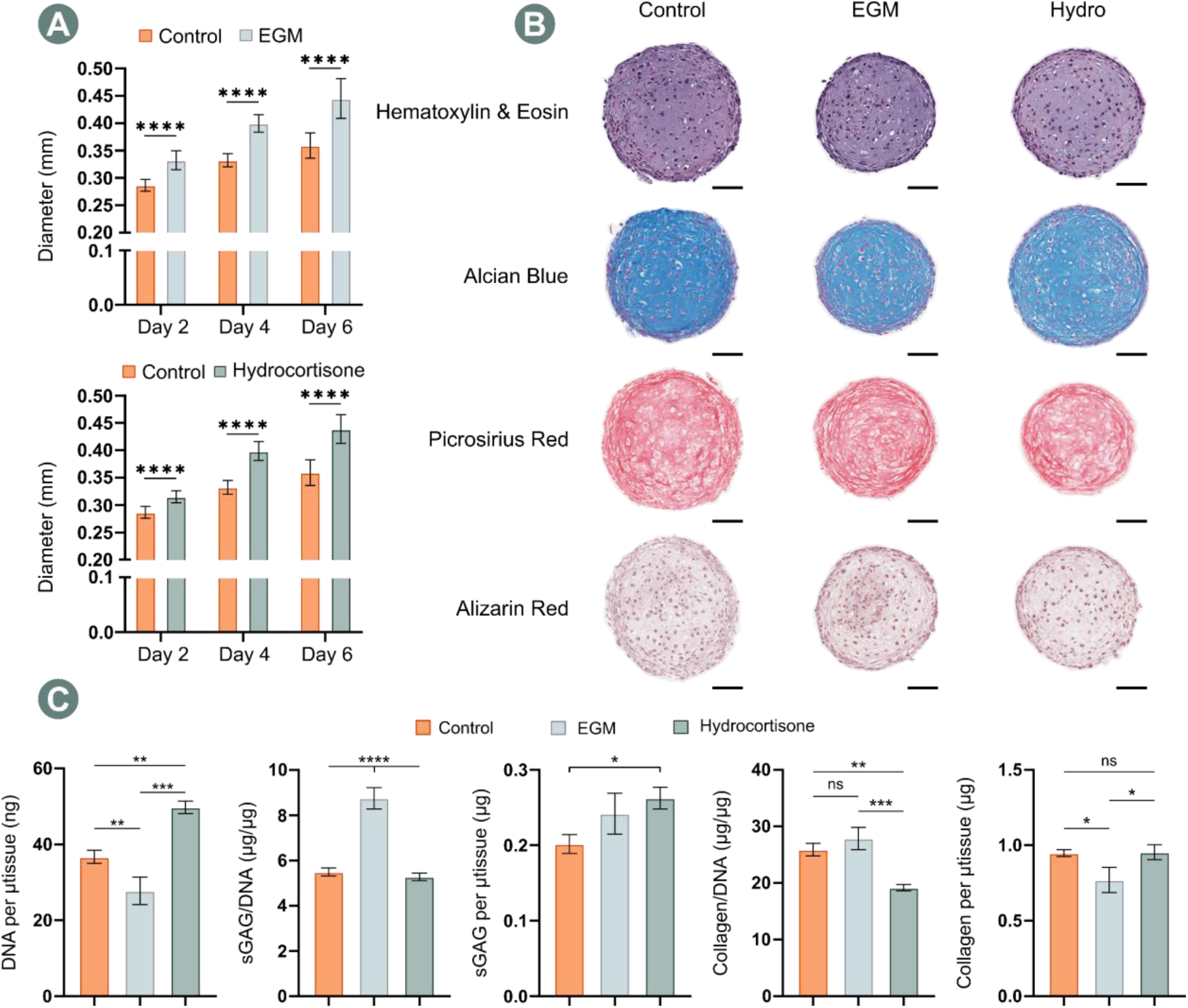
The effect of both EGM pre-soaking and hydrocortisone supplementation is lessened in a more chondrogenic hBMSC population. A) Quantification of microtissue diameter. **** denotes significance when tested using a Šídák’s multiple comparisons test, two-way ANOVA, where p < 0.0001, (N = 25, Mean ± SD). B) Histological evaluation of cartilage microtissues after 7 days of chondrogenic culture (Scale bar = 100 μm). C) Biochemical quantification of the cartilage microtissues after 21 days. * denotes significance using an Ordinary One-way ANOVA with a Tukey’s multiple comparisons test, where; ns indicates p > 0.05, * indicates p < 0.05, ** indicates p < 0.01, *** indicates p < 0.001, and **** indicates p < 0.0001 (N = 3 for control & Hydro, N = 4 for EGM, Mean ± SD).

## Discussion

This study aimed to establish a novel protocol for ensuring the generation of high-quality cartilage microtissues. Soaking hydrogel microwells in fully supplemented EGM a day prior to seeding gBMSCs was found to enhance cellular aggregation and cause a significant improvement in chondrogenesis. At day 2, all cells within the microwells of the EGM group had coalesced, forming large spherical aggregates. In contrast, a standard chondrogenic protocol yielded relatively small cell aggregates with a large number of unengaged cells surrounding the spheroids. By day 21 of culture, histological and biochemical evaluation indicated that a more cartilaginous ECM could be generated by soak loading the microwells with EGM prior to seeding. Cartilage matrix components (sGAG and collagen) were more extensively deposited in the EGM group compared to the control condition. Moreover, the increased abundance of ECM proteins was not only due to more cellular microtissues, as evident by higher DNA content after 21 days of culture, but also as a result of the increased biosynthetic output of the resident cell population within the EGM microtissues. Collectively, this preliminary study indicated that factor(s) within EGM provide potent cues capable of improving the chondrogenic capacity of an uncharacterised BMSCs population.

Next, we sought to determine if a single component within the EGM was primarily responsible for the aforementioned results using more clinically relevant hBMSCs. Within the supplement profile of EGM, basic FGF/FGF-2 (FGF) and insulin-like growth factor (IGF) were potential candidates for driving improved chondrogenesis. FGF is known to maintain MSCs in an immature state, enhance their proliferation during *in vitro* expansion and their subsequent differentiation potential [43]. Additionally, the treatment of hMSCs with FGF during expansion has given rise to enhanced chondrogenesis [44,45]. Specifically, chondrogenic aggregates formed using cells treated with FGF during monolayer expansion were larger and expressed higher proteoglycan content. Additionally, FGF-treated cells have been formed into cartilage spheroids that lacked collagen type I and expressed collagen type II in their periphery [45]. FGF signalling, although not critical for chondrogenesis, has been associated with improved chondrogenic differentiation of hMSCs [46]. However, prolonged treatment with FGF during MSC condensation and early chondrogenic differentiation has been shown to inhibit chondrogenesis, whereas the addition of other isoforms, such as FGF-9, to chondrogenic media has been shown to marginally increase matrix production during early chondrogenesis [47]. In this study, all MSCs were exposed to FGF during expansion, however exposure to FGF during the first 7 days of chondrogenic differentiation did not enhance chondrogenesis and ECM production (Supplementary Figure 2). IGF, when combined with TGF-β, is commonly discussed as a promoter of chondrogenesis in MSCs [43,48]. IGF alone has been suggested to have similar chondrogenic effects as TGF-β, stimulating proliferation, regulating apoptosis, and inducing the expression of chondrogenic markers in BMSCs. Moreover, the two growth-factors have demonstrated additive effects, resulting in gene expression analogous to human primary culture chondrocytes [49]. We failed to see a similar results in this study, as there was no discernible benefit associated with supplementing CDM with IGF (Supplementary Figure 2).

The growth factor hydrocortisone emerged as the principle driving factor within the EGM supplement capable of promoting more robust chondrogenesis. Intra-articular injection of glucocorticoids, such as hydrocortisone, is a longstanding means of managing arthritis. Primarily, glucocorticoid therapy aims to provide symptomatic relief, reducing inflammation and pain within an affected joint. However, the use of such steroidal agents has been discouraged for the treatment of OA due to their undesirable effects on cartilage metabolism [50]. Despite this, chondroprotective properties and other putative benefits of glucocorticoid treatment have been suggested. More recently, the chondroprotective capacity of hydrocortisone has been found to be heavily dose-dependent, with beneficial changes associated with low doses both *in vitro* and *in vivo*, whereas higher doses result in deleterious effects [51]. *In vitro*, the exposure of MSCs to synthetic glucocorticoids for the initiation of chondrogenesis has been well established through the use of dexamethasone (DEX) [52,53]. The role of DEX in promoting chondrogenesis has been elucidated through studies demonstrating that glucocorticoids directly regulate the expression of cartilage ECM genes and/or enhance TGF-β-mediated effects on their expression. Specifically, a positive interaction between TGF-β and glucocorticoid signalling pathways, which are mediated by the glucocorticoid receptor, have been demonstrated during chondrogenesis [54]. The impact of hydrocortisone, an adrenocortical hormone, is much less documented. It has been reported to be found in FBS, where it is important in the modulation of MSC functions such as growth and adhesion [55]. As such, it is often included in serum-free medium formulations. Additionally, hydrocortisone has been used to ‘activate’ multipotent MSCs for adipogenic differentiation, while its addition during passaging helps to preserve the self-maintenance capacity of MSCs [56]. In the context of chondrogenesis, hydrocortisone supplementation in 3D culture with human chondrocytes has been shown to optimise ECM metabolism. In particular, exposure to physiological levels of hydrocortisone was linked with an enhanced capacity to synthesise ECM components (aggrecan, collagen type II, and fibronectin) whilst decreasing the activity of catabolic pathways (suppression of the IL1 catabolic pathway - reduced intracellular IL1-α and −β as well as IL1RI) [57]. Collectively, these results indicate that corticosteroids can be beneficially leveraged in the engineering of high-quality cartilage microtissues using MSCs.

Whilst the mechanism of action remains unclear, we present evidence that suggests EGM/hydrocortisone treatment can improve the chondrogenic potential of heterogeneous MSC populations. Investigation into the full effects of this novel chondrogenic protocol in terms of its regulation/re-activation of MSC subpopulations, and/or its effect on chondrogenic genes that are co-regulated *via* glucocorticoid receptors would provide interesting additional insight and could help design enhanced chondrogenic cultures in the future. Given the evidence that selection of superior chondrogenic donors *in vitro* can translate into improved *in vivo* outcomes [58,59], the data presented in this work represents a simple alternative method for maximising the chondrogenic capacity of MSC populations that exhibit inherently limited chondrogenesis. As such, effectively implementing this novel protocol can result in the formation of high-quality cartilage microtissues. To this end, our preliminary data (Supplementary Figure 3) suggests that exposure to hydrocortisone at a concentration of 0.2 μg/mL may be beneficial for chondrogenic culture. This evidence, coupled with our findings relating to the potency of EGM soak loading indicates that a similar soaking, or short-term exposure strategies (<7 days) may be the most effective means of implementing hydrocortisone treatment within a chondrogenic culture regime. Ultimately, this work provides a platform to generate larger engineered cartilage through self-organisation of these high-quality building blocks without the need for additional cells, unfeasible numbers of microtissue units, or compromising the quality of the final construct.

## Conclusion

Collectively, the results of this study indicate that pre-treatment *via* EGM pre-soaking or the supplementation of chondrogenic differentiation medium with hydrocortisone can provide a simple and potent means of improving chondrogenesis in heterogeneous MSC cohorts. This work could enable the generation of more scalable engineered cartilages by ensuring the formation of high-quality cartilage microtissue building blocks without the need for extensive cell immunophenotyping.

## Acknowledgements

This publication was developed with the financial support of Science Foundation Ireland (SFI) under grant number 12/RC/2278 and 17/SP/4721. This research is co-funded by the European Regional Development Fund and SFI under Ireland’s European Structural and Investment Fund. This research has been co-funded by Johnson & Johnson 3D Printing Innovation & Customer Solutions, Johnson & Johnson Services Inc.

