## Supporting information for "Engineering High-Quality Cartilage Microtissues using Hydrocortisone Functionalised Microwells"

### Supplementary Information

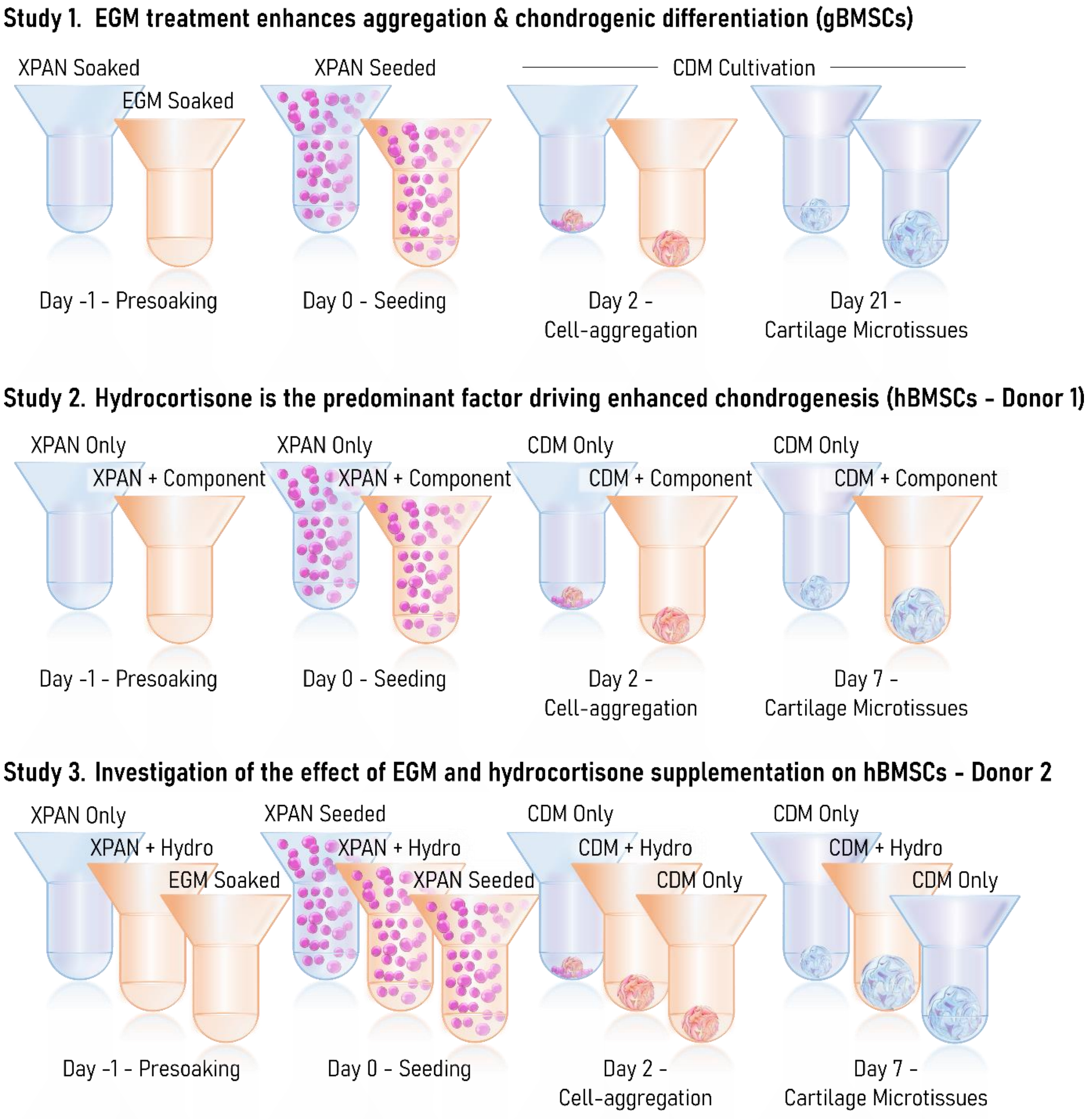


***Supplementary Figure 1.*** *Study Schematic. In study 1) Microwells are soak loaded with EGM, after which a normal chondrogenic protocol is undertaken. Study 2) Factors are soak loaded into the microwells and added to culture medium for the entire culture period. Study 3) Hydrocortisone is compared to EGM soak loading only and traditional chondrogenic cultivation in a second human donor. Both studies 2 & 3 are undertaken over 7 days to study the effect of treatment on initial aggregation and early chondrogenic differentiation of human mesenchymal stem/stromal cells (hMSCs).*

14

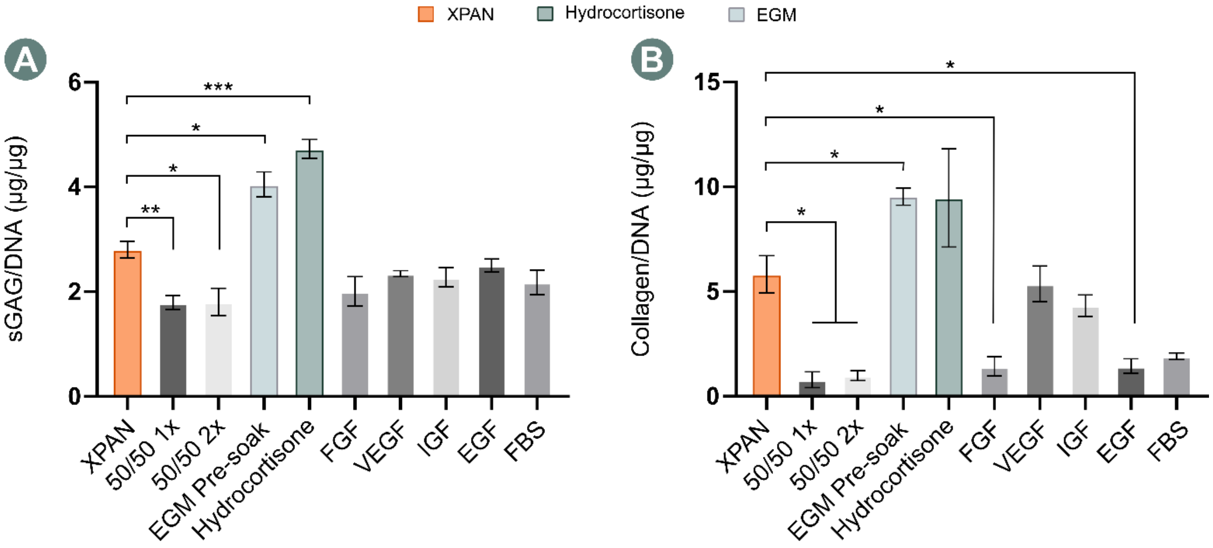


***Supplementary Figure 2.*** *Determining the driving factor in EGM. Biochemical quantification after 7 days of chondrogenic culture of A) sGAG normalised to DNA, (N = 3, Mean ± SD) and B) collagen normalised to DNA, (N = 3, Mean ± SD). * denotes significance using a Brown-Forsythe and Welch One-way ANOVA, where * indicates p < 0.05, ** indicates p < 0.01, and *** indicates p < 0.001*

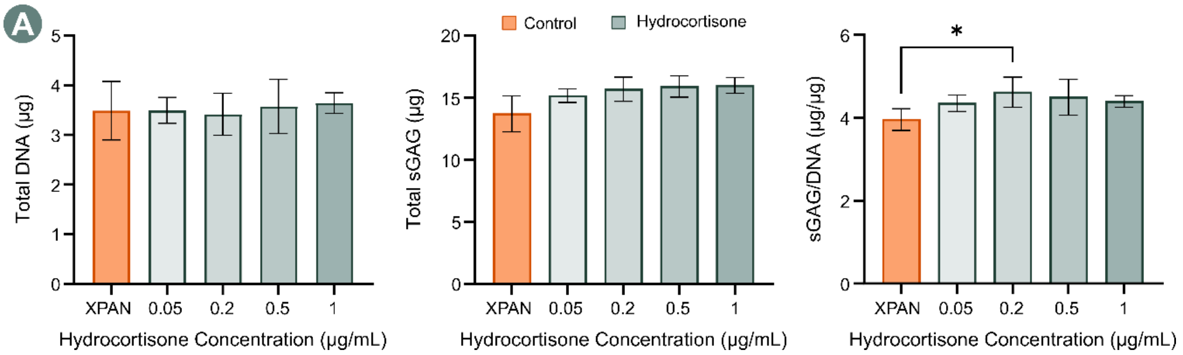


***Supplementary Figure 3.*** *A) Biochemical quantification of the cartilage microtissues after 7 days of chondrogenic culture. Identifying 0.2 µg/mL as an effective concentration of hydrocortisone for generating significantly higher levels of sGAG/DNA. * denotes significance using a two-tailed, unpaired Welch's t-test, where p < 0.05 (N = 4, Mean ± SD).*

| Experimental Group/Media type | FBS | FGF-2 | Hydro | VEGF | IGF | EGF | TGF-β | ITS | Dex | AA |
|---|---|---|---|---|---|---|---|---|---|---|
| **XPAN** | Yes (10 %v/v) | Yes (5 ng/mL) | No | No | No | No | No | No | No | No |
| **EGM (%v/v)** | Yes (5%) | Yes (0.4%) | Yes (0.04%) | Yes (0.1%) | Yes (0.1%) | Yes (0.1%) | No | No | No | Yes (0.1%) |
| **CDM** | No | No | No | No | No | No | Yes (10 ng/mL) | 1× | Yes (100 nM) | Yes (50 µg/mL) |
| **50/50 1×** | • Yes (7.5% - XPAN/EGM<br>• Yes (2%) - CDM/EGM) | • Yes (2.5 ng/mL + 0.2%) - XPAN/EGM<br>• Yes (0.2%) - CDM/EGM) | Yes (0.02%) | Yes (0.05%) | Yes (0.05%) | Yes (0.05%) | • No – XPAN/EGM<br>• Yes (10 ng/mL) – CDM/EGM | • No – XPAN/EGM<br>• Yes (1×) – CDM/EGM | • No – XPAN/EGM<br>• Yes (100 nM) – CDM/EGM | • Yes (0.05%) – XPAN/EGM<br>• Yes (0.05% + 50 µg/mL) – CDM/EGM |
| **50/50 2×** | • Yes (7.5%) XPAN/EGM<br>• Yes (5%) - CDM/EGM) | • Yes (2.5 ng/mL + 0.4%) - XPAN/EGM<br>• Yes (0.4%) - CDM/EGM | Yes (0.04%) | Yes (0.1%) | Yes (0.1%) | Yes (0.1%) | • No – XPAN/EGM<br>• Yes (10 ng/mL) – CDM/EGM | • No – XPAN/EGM<br>• Yes (1×) – CDM/EGM | • No – XPAN/EGM<br>• Yes (100 nM) – CDM/EGM | • Yes (0.1%) – XPAN/EGM<br>• Yes (0.1% + 50 µg/mL) – CDM/EGM |
| **Hydro** | • Yes (10%) XPAN/Hydro<br>• No - CDM/Hydro | • Yes (5 ng/mL) – XPAN/Hydro<br>• No – CDM/Hydro | Yes (0.04%) | No | No | No | • No – XPAN/Hydro<br>• Yes (10 ng/mL) – CDM/Hydro | • No – XPAN/EGM<br>• Yes (1×) – CDM/Hydro | • No – XPAN/EGM<br>• Yes (100 nM) – CDM/Hydro | • No – XPAN/Hydro<br>• Yes (50 µg/mL) – CDM/Hydro |
| **FGF** | • Yes (10%) XPAN/FGF<br>• No - CDM/FGF | • Yes (5 ng/mL) – XPAN/FGF<br>• Yes (0.4%) – CDM/FGF | No | No | No | No | • No – XPAN/FGF<br>• Yes (10 ng/mL) – CDM/FGF | • No – XPAN/FGF<br>• Yes (1×) – CDM/FGF | • No – XPAN/FGF<br>• Yes (100 nM) – CDM/FGF | • No – XPAN/FGF<br>• Yes (50 µg/mL) – CDM/FGF |

| | | | | | | | | | | |
|---|---|---|---|---|---|---|---|---|---|---|
| **VEGF** | • Yes (10%) XPAN/VEGF<br>• No - CDM/VEGF | • Yes (5 ng/mL) – XPAN/VEGF<br>• No – CDM/VEGF | No | Yes (0.1%) | No | No | • No – XPAN/VEGF<br>• Yes (10 ng/mL) – CDM/VEGF | • No – XPAN/VEGF<br>• Yes (1×) – CDM/VEGF | • No – XPAN/VEGF<br>• Yes (100 nM) – CDM/VEGF | • No – XPAN/VEGF<br>• Yes (50 μg/mL) – CDM/VEGF |
| **IGF** | • Yes (10%) XPAN/IGF<br>• No -CDM/IGF | • Yes (5 ng/mL) – XPAN/IGF<br>• No – CDM/IGF | No | No | Yes (0.1%) | No | • No – XPAN/IGF<br>• Yes (10 ng/mL) – CDM/IGF | • No – XPAN/IGF<br>• Yes (1×) – CDM/IGF | • No – XPAN/IGF<br>• Yes (100 nM) – CDM/IGF | • No – XPAN/IGF<br>• Yes (50 μg/mL) – CDM/IGF |
| **EGF** | • Yes (10%) XPAN/EGF<br>• No - CDM/EGF | • Yes (5 ng/mL) – XPAN/EGF<br>• No – CDM/EGF | No | No | No | Yes (0.1%) | • No – XPAN/EGF<br>• Yes (10 ng/mL) – CDM/EGF | • No – XPAN/EGF<br>• Yes (1×) – CDM/EGF | • No – XPAN/EGF<br>• Yes (100 nM) – CDM/EGF | • No – XPAN/EGF<br>• Yes (50 μg/mL) – CDM/EGF |
| **FBS** | • Yes (10%) XPAN/FBS<br>• Yes (5%) - CDM/FBS | • Yes (5 ng/mL) – XPAN/FBS<br>• No – CDM/FBS | No | No | No | No | • No – XPAN/FBS<br>• Yes (10 ng/mL) – CDM/FBS | • No – XPAN/FBS<br>• Yes (1×) – CDM/FBS | • No – XPAN/FBS<br>• Yes (100 nM) – CDM/FBS | • No – XPAN/FBS<br>• Yes (50 μg/mL) – CDM/FBS |

***Supplementary Table 1.*** *Experimental groups and corresponding media formulations*

20
